# An improved SIFT algorithm based method for detecting of tumor tissue movement in respiratory motion

**DOI:** 10.1101/2025.08.05.668829

**Authors:** Qingya Pan, Shengjie Luan, Fuke Zhang, Linzhao Tian, Guishu Wu, Yizhong Fan

**Affiliations:** Department of ChemoRadiotherapy Oncology, The Center Hospital of Qinghe,HeBei,China

**Keywords:** SIFT, cross- correlation, moved phantom, rotated phantom

## Abstract

**Background:** This paper intends to make use of the excellent performance of the SIFT algorithm to accurately calculate the displacement of tumor tissues during respiratory movement.

**Material and method:** First, we moved phantoms to up,down,right,left respectively to simulate the different directions of movement during respiratory. And rotated phantom to left and right respectively by the angle of 5,10,30,45. Then, used SIFT and cross-correlation algorithms to search the phantoms. Last, use the t-test to check the differences between different directions.

**Resul:** The t-test result proved that there was no significant difference in the detection of moving phantom in all directions. There was no statistical difference between the SIFT of the moving motif and the cross-correlation. The detection results of rotating phantom indicated that the SIFT test results could be used to detect distortion.

**Conclusion:** SIFT algorithm can be used to calculate the distortion caused by the thoracic tissue during respiratory movement.

## 1. Introduction

Lung cancer is a disease with high mortality rate. Nearly 70% of cancer patients need to receive radiotherapy during the course of treatment.^1^ However, determining the accuracy of tumor movement during the breathing process is a challenge. If the movement of tumors during the breathing process can be accurately grasped, the radiation exposure endured by patients can be significantly reduced. So far, 4DCT is mostly used in clinical to determine the movement of tumors, which is time-consuming and labor-intensive.^2^

Previously, many people have made attempts to improve the accuracy of automatic detection. Emilie E. proposed a dynamic tumor tracking is a motion management technique where the radiation beam follows a moving tumor in real time.^3^ Christoph Hoog Antink use a high-sampling rate terahertz (THz) homodyne spectroscopy system to estimate thoracic movement from healthy subjects performing breathing at different frequencies^4^. Evan SilversteinIn adapted the Microsoft Kinect v2 sensor to trace and record a patient’s respiratory motion.^5^ Mari Honda used a breathing movement sensor for chest radiography during inspiration in children aged less than 3 years.^6^ Melanie Habatsch, using free-breathing, self-gating 4D magnetic resonance imaging workflow to assess movement of breast and organ-at-risks.^7^

These studies have greatly improved the accuracy of tumor treatment and significantly reduced the damage to the surrounding organs at risk.Scale-Invariant Feature Transform features (SIFT) are widely used in applications for target detection^8^, tracking^9^, classification^10^, image matching^11^, and constructing mosaic images^12^. Due to its invariance to rotation, and translation, local invariant features are expected to be feasible features with defective image. However, the movement of the chest tissue is not a simple shift, and there is a certain degree of twisting with breathing. This paper intends to make use of the excellent performance of the SIFT algorithm to accurately calculate the displacement of tumor tissues during respiratory movement.

## 2. Materials and methods

The thoracic phantom employed in this study is showed in Figure 1, along with the synthetic nodules were attached before CT imaging. The phantom does not contain lung parenchyma, so the space within the vascular structure is filled with air. The phantom was scanned using a Philips 16-row scanner (Mx8000 IDT, Philips Healthcare, Andover, MA). Scans were acquired with, pitch (100), and slice (3.0mm) were reconstructed slice thicknesses (3.0mm) and reconstruction kernels (1.5mm). The phantom position was re-positioned between different nodule layouts. Images in this paper are 512*512 for each CT slice. We simulated changes in tumor position with respiratory movement by moving the position of synthetic nodules showed in figure2, and rotation of tumor position in figure3.

**Figure 1.**
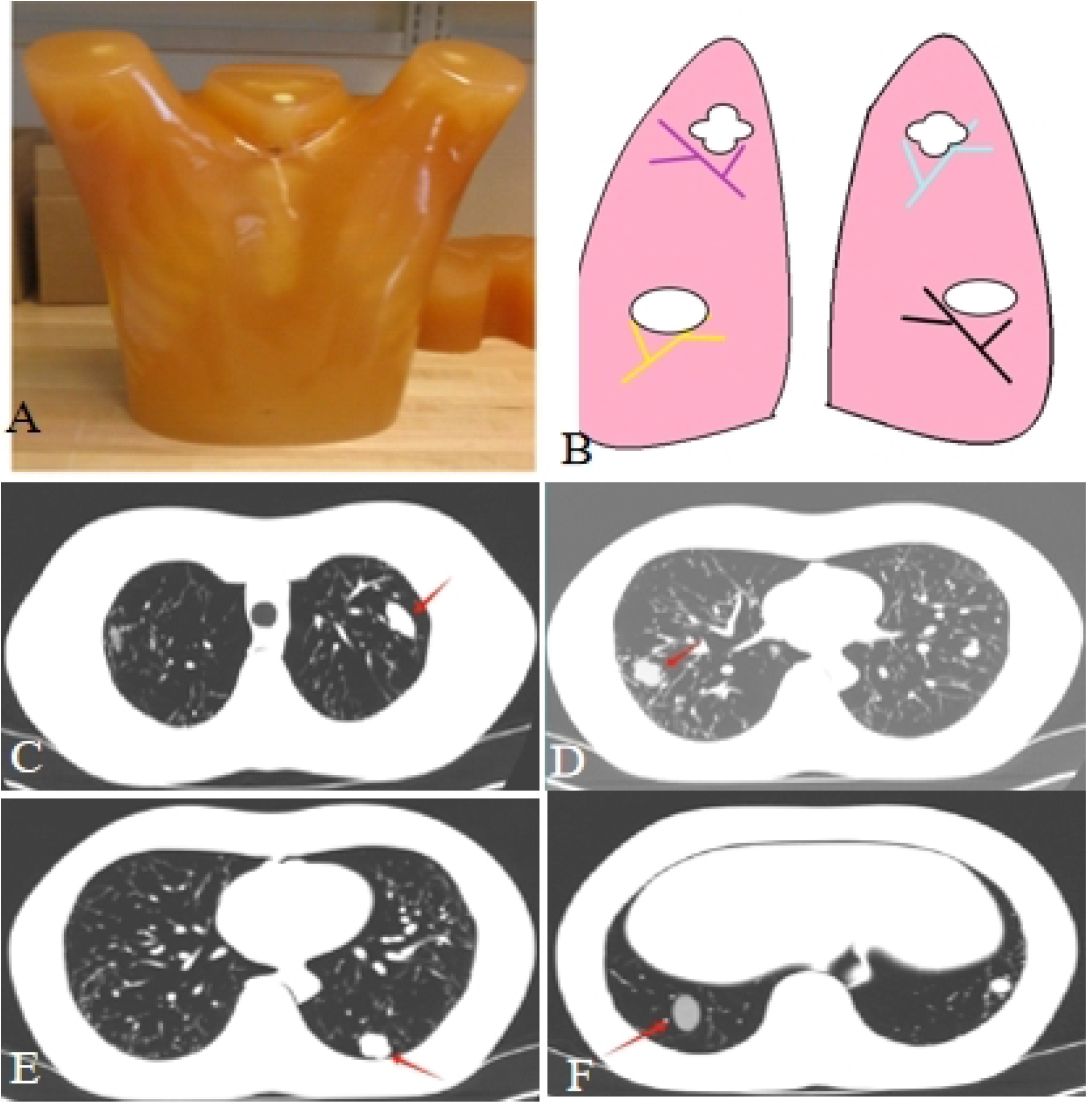
The thoracic phantom.

### 2.1 Scale-Invariant Feature Transform (SIFT)

Scale-Invariant Feature Transform (SIFT) descriptor is based on gradient orientation histograms computed in the region around feature points. The performance of target detection can be improved by optimizing the SIFT parameters for specific scenarios.

#### A. Construct Gaussian space

The difference of the Gaussian scale-space is computed to obtain SIFT

key points as below:

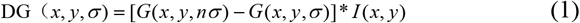

Here, x and y are the pixel coordinates of the processed image *I* (*x, y*), *G*(*x, y,σ*) is the Gauss filter having variable scale, *n* is the scale multiplication coefficient, and D*G*(*x, y,σ*) is the Gauss difference image.

In this space, scale-space extremas are detected within the neighborhood of each point in the previous and next scale-space image.

#### B. Eliminate key-points

Key-points go through a further elimination process by using a cornerness measure. For this purpose,Hessian matrix is computed for each key-point

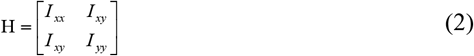

Here, 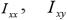 are the directional derivatives computed in the horizontal, diagonal, and vertical directions, respectively. For the keypoint located on a corner, both eigenvalues of the corresponding Hessian matrix are distinct and take high values.

Equation (3) meets these constraints and is used to decide whether a keypoint is located on a corner or not:

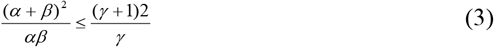

In the equation, *α* and *β* are the eigenvalues of the Hessian matrix. The corner coefficient *γ*is a parameter to be considered in feature extraction.

#### C. Orientation Assignment and Descriptor Definition

An orientation is computed and assigned for each key-point to achieve rotation in-variance. A gradient histogram is constructed, and the local peak within the 80% of the highest peak is assigned as the orientation of the key-point. If more than one local peaks exist, then a key-point is defined for each orientation.

A SIFT descriptor is constructed using the gradient magnitudes and orientations around the key-point detected before. The scale of the key-point is used to create a Gaussian window and to weight the gradient magnitudes around the key-point with this window. Gradient orientations are rotated according to the predefined key-point orientation. Finally, gradient histograms are constructed for the four by four regions around the key-point and each histogram is inserted into a row vector in order to construct the SIFT descriptors.

#### D. Feature Matching

The Euclidian distance measure is used to match these descriptors. First, the distances between all descriptors are found. Then, for each descriptor, find the two closest descriptors.

In this paper, We manually delineate the edge point of the known synthetic nodules, and then directly take the points on the mold contour as the screened extreme points to calculate the descriptor and feature match.

### 2.2 Cross-correlation

The cross-correlation function is a measure of the similarity between two images.It calculates the cross-correlation in the spatial or frequency domain to determine the movement of the image in the spatial domain. the native MATLAB functions xcorr2 compute the cross-correlation functions by the dot-product method (eq 4), where the inputs are matrices, respectively, for 2D correlation.

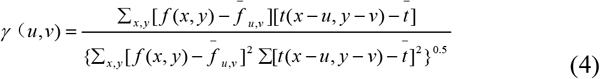

Where f is original image,t is template 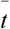 is mean of the template 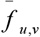 is mean of the template of *f* (*x, y*).

In this experiment, we adopted a total of eight synthetic nodules with different shapes and gray values.The nodules are: 1 spherical,HU 100, 2 spherical, HU 100, 3 spherical,HU -630, 4 spherical, HU -630, 5 elliptical,HU 100, 6 lobulated,HU 100,7 spiculated,HU -630, 8 lobulated, HU -630. We moved these nodules up, down, left and right respectively,and obtained moved nodules are showed in Table1.

**Table 1.** List of moved nodules 1.

| Noddles | Position | Moved Noddles |  |  |  |
| --- | --- | --- | --- | --- | --- |
|  |  | Up | Down | Right | Left |
| 1 |  | 1_up | 1_down | 1_right | 1_left |
| 2 |  | 2_up | 2_down | 2_right | 2_left |
| 3 |  | 3_up | 3_down | 3_right | 3_left |
| 4 |  | 4_up | 4_down | 4_right | 4_left |
| 5 |  | 5_up | 5_down | 5_right | 5_left |
| 6 |  | 6_up | 6_down | 6_right | 6_left |
| 7 |  | 7_up | 7_down | 7_right | 7_left |
| 8 |  | 8_up | 8_down | 8_right | 8_left |

Since all the nodules of 1-4 are spherical, there is no difference before and after rotation. The image of moved nodules are showed in Figure 2. We rotated nodules 5-8. The rotated nodules images are showed in Figure 2. The obtained rotated nodules showed in Table2. The SIFT algorithm and the cross-correlation algorithm were respectively used to detect these nodules.The programs used in this paper were developed using the MATLAB 2024a (The MathWorks, Inc, Natick, MA).

**Table 2.** List of rotated nodules 1.

|  |  | Rotated Noddles Position |  |  |  |
| --- | --- | --- | --- | --- | --- |
| Rotation angle |  | L_up | L_down | R_up | R_down |
| L | 5 | 5_L5 | 6_L5 | 7_L5 | 8_L5 |
|  | 10 | 5_L10 | 6_L10 | 7_L10 | 8_L10 |
|  | 30 | 5_L30 | 6_L30 | 7_L30 | 8_L30 |
|  | 45 | 5_L45 | 6_L45 | 7_L45 | 8_L45 |
| R | 5 | 5_R5 | 6_R5 | 7_R5 | 8_R5 |
|  | 10 | 5_R10 | 6_R10 | 7_R10 | 8_R10 |
|  | 30 | 5_R30 | 6_R30 | 7_R30 | 8_R30 |
|  | 45 | 5_R45 | 6_R45 | 7_R45 | 8_R45 |

**Figure 2.**
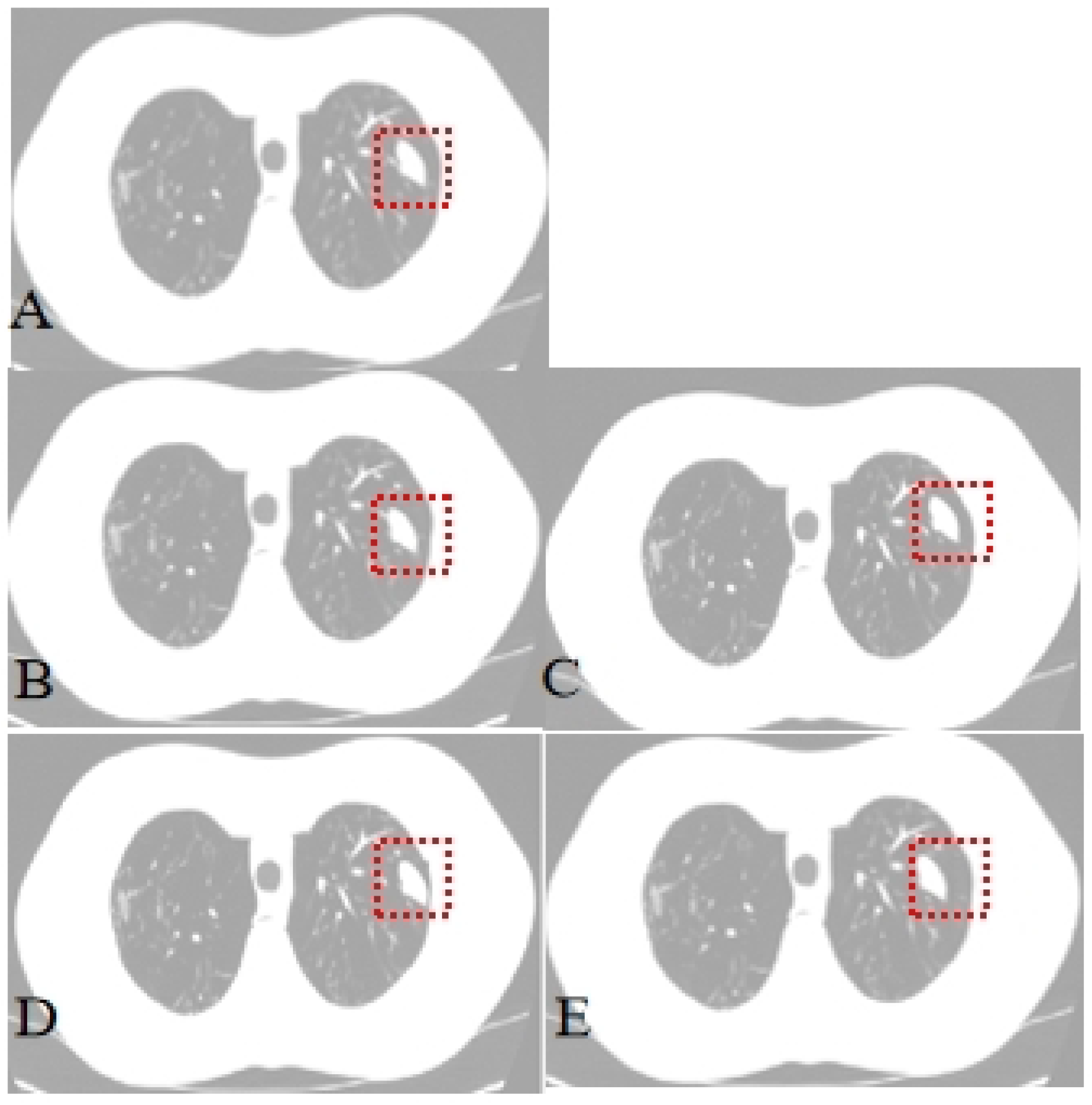
shows the move.

**Figure 3.**
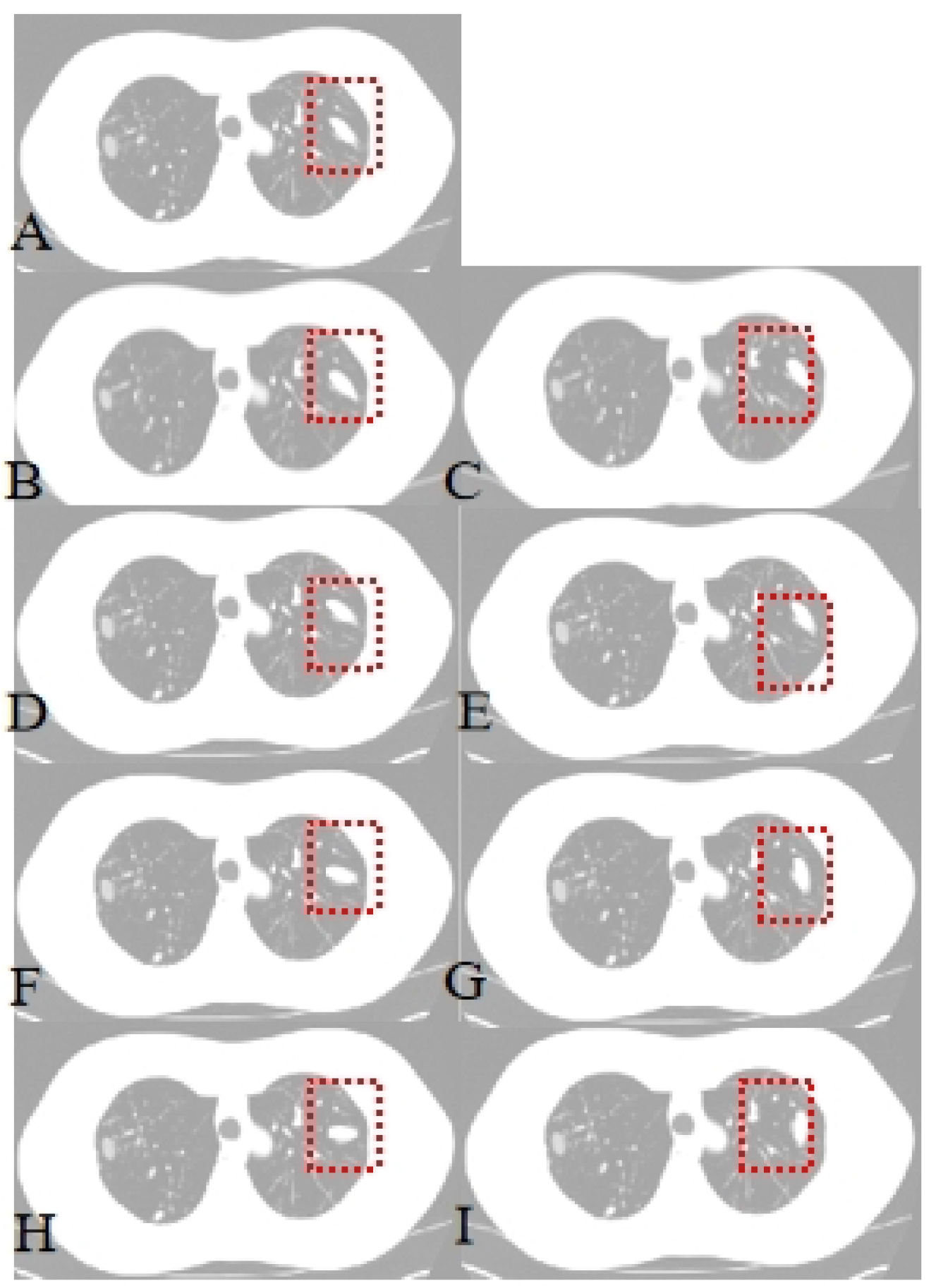
shows the ro.

**Figure 4.**
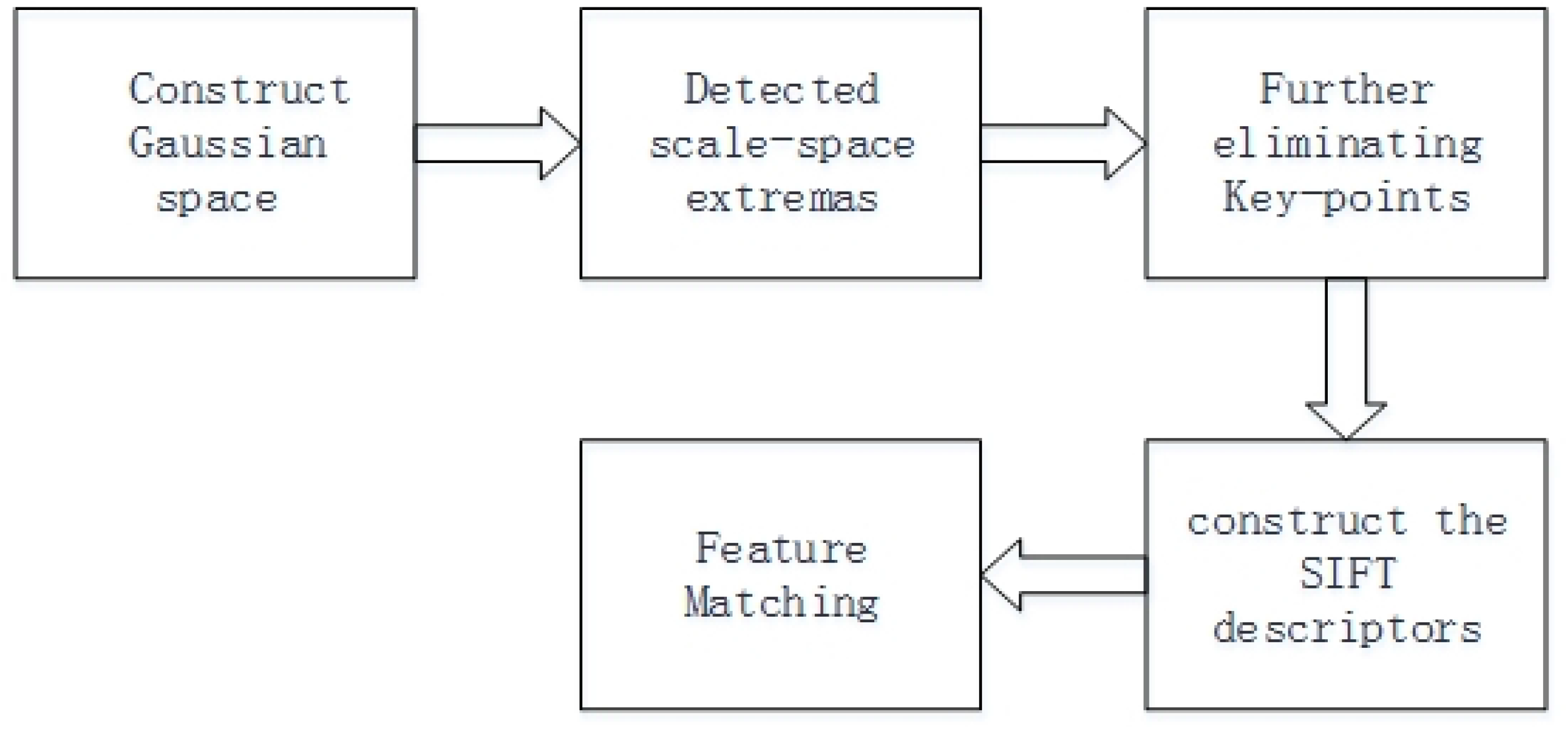
flow chart of SIFT algorithm.

**Figure 5.**
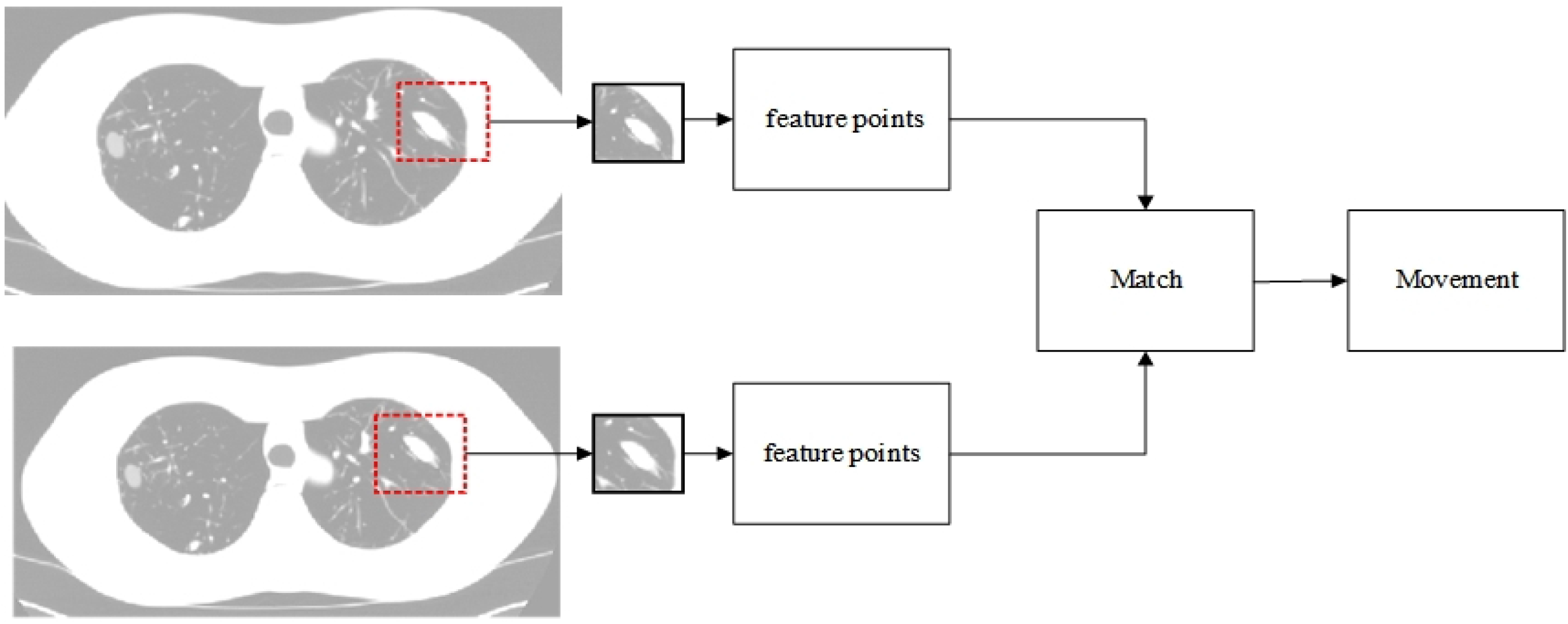
flow chart of SIFT algorithm we used.

## 3. Result

The statistical results of this experiment are shown in Table3. Table 3 shows the influence of different movement directions on the search results of the correlation and SIFT algorithms. Most t-test results in Table3 showed P<0.05. Indicating that there was no statistical significance among the detection results of moving nodules in different directions. T-test for nodules shows obvious difference both SIFT and cross-correlation algorithms (nodules5, right, P=0.128; nodules6 down, P=0.312; nodules7, left, P=0.534). Indicating that there was statistical significance among the detection results of these moving nodules. Figure 6 presents a comparison between the SIFT and cross-correlation detection results across eight synthetic nodules. Figure 7 shows Box plot of the detection results of the rotating nodules at various angular positions. The nodules are :(nodules 5 elliptical,HU 100), (nodulee6 lobulated,HU 100),(nodule 7 spiculated,HU -630), (nodule 8 lobulated, HU -630).

**Table 3.** Result of t test for moved noddle 1.

| Noddles | Correlation |  |  | SIFT |  |  |
| --- | --- | --- | --- | --- | --- | --- |
|  | down | right | left | down | right | left |
| 1 | 0.02 | 0.00 | 0.02 | 0.01 | 0.03 | 0.04 |
| 2 | 0.01 | 0.01 | 0.00 | 0.03 | 0.02 | 0.01 |
| 3 | 0.03 | 0.02 | 0.01 | 0.04 | 0.05 | 0.02 |
| 4 | 0.01 | 0.02 | 0.04 | 0.03 | 0.04 | 0.03 |
| 5 | 0.02 | 0.128 | 0.01 | 0.00 | 0.214 | 0.01 |
| 6 | 0.312 | 0.03 | 0.03 | 0.302 | 0.01 | 0.03 |
| 7 | 0.03 | 0.02 | 0.534 | 0.02 | 0.05 | 0.749 |
| 8 | 0.02 | 0.03 | 0.04 | 0.00 | 0.01 | 0.02 |

**Figure 6.**
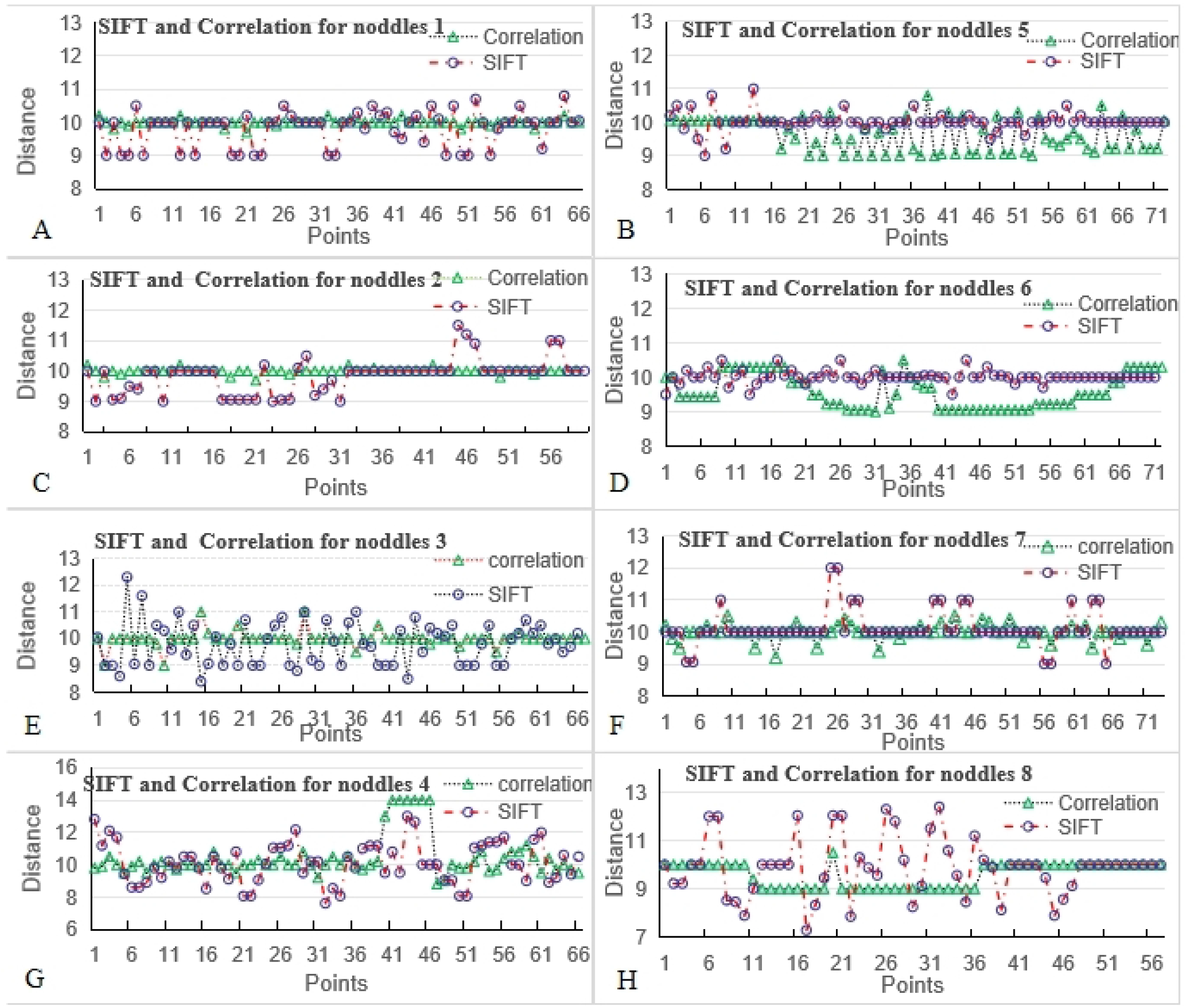
presents a comparison between the SIFT.

**Figure 7.**
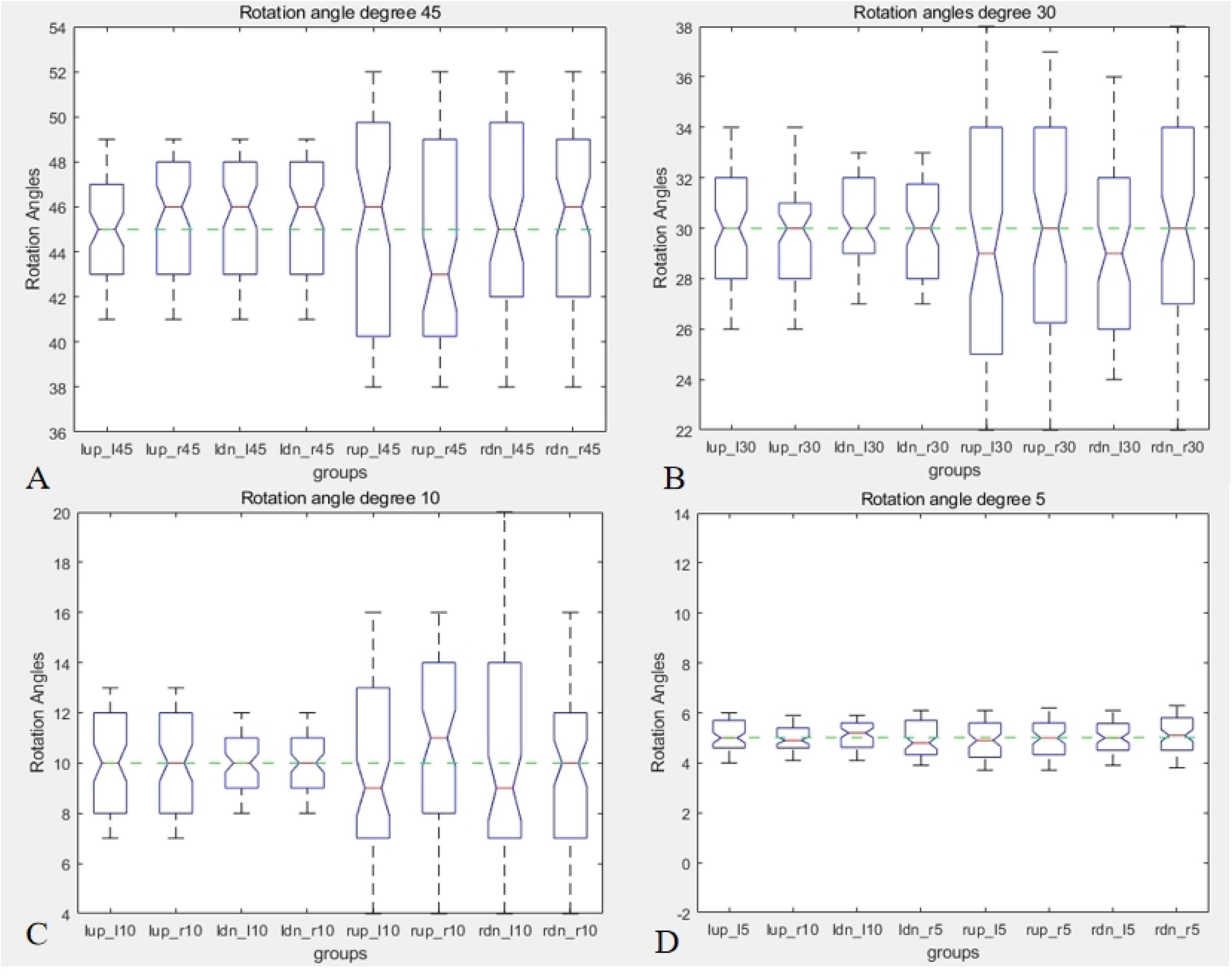
The detection results of the rotating nodule at various.

## 4. Discussion

The SIFT algorithm has been applied in many fields, and many people have made attempts achieved significant results^15-20^.

In this paper, we attempt to use the SIFT algorithm to calculate the movement of chest tissue during respiratory movement. In previous methods, the calculation of movement changes did not take into account rotational distortion. We attempt to use the SIFT algorithm to determine the distorted changes in the chest tissue during respiratory.

Table3 shows the results of the t-test under different movement directions. We used the moving nodules in the down, right and left directions to compare with the upward moving nodules. It can be seen from Table3 that the movement of the nodules in each direction at each position has no statistical significance on the detection results. The detection results are significantly different from those in several other directions. We observed the position of the noddle in the phantom and found that during the movement, the moved noddle crossed with the ribs, and the boundary of the noddle was severely blurred, which led to inaccurate detection results.

As can be seen in Figure6, under the premise of a moving distance of 10, the detection results of each model all fluctuate around 10. As shown in Figure6, the fluctuation of nodules 1,2,5,6 are significantly smaller than that of nodules 3,4,7,8, indicating that nodules 1,2,5,6 are more accurate than other moved nodules. This is because their gray values are relatively high (HU 100), resulting in a higher contrast with the surrounding tissues. During our calculation process, we need to perform the conversion to grayscale image, which results in a significant difference in their gray values. The cross-correlation detection results of each module in nodules1-4 are also better than those in nodules 5-8. This is because the nodules in the nodules 1-4 are all circular, and their boundaries change less during the movement process, making the detection results more accurate.

The results detected by the SIFT algorithm fluctuate more compared to cross-correlation. This is because we omitted the key point filtering process in order to accurately determine the positions of key points. The calculation of all key points is to achieve the known edge of the selected synthetic nodule. In Figure7, the SIFT algorithm can calculate the rotation angle of the nodules 5-8. In right side synthetic nodules (such as:nodules 7,nodules 8) with relatively low gray values (HU -630), the calculation results fluctuate more. The fluctuation of left side synthetic nodules are relatively small, which is also due to the gray value (HU 100). Perhaps improvements can be made in this aspect in future calculations.

In addition, when the rotation angle is small, the error is also relatively small. This is mainly because the rotation angle affects the change of the gray value around the nodules. The larger the rotation angle, lead to the greater the change around nodules, and the greater the probability of error during search.

Another important point is that neither the SIFT algorithm nor the cross-correlation algorithm can provide satisfactory detection results for uniform media. The search in these places still requires further research on relevant algorithms.

## 5. Conclusion

This paper calculated the movement and rotation of the model, confirms that the SIFT algorithm can be used to calculate the distortion caused by the thoracic tissue during respiratory movement.The accuracy of the rotation Angle still needs to be further improved.

## Acknowledgments

This study is supported by the key science and technology research plan of HeBei Provincial Health Commission Scientific Research Fund Project: 20201329. This study is approved by the ethics committee of Qinghe Central Hospital.

